# Early regional lymph node activation drives influenza vaccine responses in an ancestrally diverse cohort

**DOI:** 10.1101/2024.10.28.620725

**Authors:** Jacqueline HY Siu, Sofia Coelho, Aime Palomeras, Sandra Belij-Rammerstorfer, Chloe H Lee, Tamara Strobel, Christopher Thorpe, Charandeep Kaur, Tom Cole, Nico Remmert, Jamie Fowler, Sam Pledger, Kyla B Dooley, Daniel Opoka, Tamas Szommer, Samantha Vanderslott, Pontiano Kaleebu, Anita Milicic, Donald B Palmer, Teresa Lambe, Brian D Marsden, Hashem Koohey, Mark Coles, Calliope A Dendrou, Katrina M Pollock

## Abstract

Early *in vivo* dynamics of human immune-cell activation across regionally activated lymphoid tissue sites upon immunisation are poorly characterised in ancestrally diverse individuals. Here, we profiled draining and non-draining axillary lymph nodes (dLNs and ndLNs) by ultrasound-guided fine-needle aspiration (FNA) in 13 Black and Asian ancestry individuals, before and 3-7 days after vaccination with adjuvanted influenza vaccine. Draining but not ndLNs rapidly increased in size post-vaccination, by day 3, with distinct cellular dynamics determined through single cell multiomics. Dissecting LN cellular diversity into 42 lymphoid and non-lymphoid cell states, post-vaccination cell abundance changes were observed across all LNs, but dLNs were specifically characterised by CD4^+^ T follicular helper (CD4^+^ Tfh) cell expansion. Gene expression analysis revealed a dLN post-vaccination hub of multicellular activity defined by CD4^+^ Tfh signalling, cross-compartmental activation, translation, and enhanced antigen-presentation capacity. Thus, robust responses to intramuscular immunisation transcending ancestral inter-individual variation are elicited through temporal, anatomical and cellular lymphatic co-ordination with implications for vaccine design in ancestrally diverse populations.

**Summary:** In this study of ancestrally diverse young adults, the temporarily co-ordinated response to an adjuvanted influenza vaccine at lymph nodes local to (draining) and distal from (non-draining) the injection site, reveals early regulation of cellular kinetics and anatomical hierarchy of the innate and adaptive immune responses.

## Introduction

Within lymph nodes (LNs), germinal centre (GC) interactions between B cells and CD4^+^ T follicular helper (CD4^+^ Tfh) cells drive the high-affinity antibody responses typically required for vaccine-dependent, long-term protection against pathogens. Studies in animal models suggest that early LN immune responses dictate vaccine priming efficacy (Liang et al., 2017), but in humans the early *in vivo* dynamics of immune-cell activation in secondary lymphoid tissues upon immunisation remain poorly characterised. Instead, human research in vaccination relies on peripheral blood samples, and whilst the frequency and activation of circulating CD4^+^ Tfh cells and plasmablasts increase in young adults within the first week post-immunisation with recall antigens, the blood does not contain GCs (Vella et al., 2019). LN fine-needle aspiration (FNA) is emerging as a tractable methodology to address unexplored aspects of human lymphatic biology *in vivo* (Havenar-Daughton et al., 2020). Recent studies employing ultrasound-guided FNA to sample draining LNs (dLNs) have utilised FNA-derived dLN cell profiling to demonstrate the evolution of B and CD4^+^ Tfh cell responses to non-adjuvanted influenza and COVID-19 mRNA vaccination weeks and even months after immunisation (Turner et al., 2020; Turner et al., 2021; Mudd et al., 2022; Schattgen et al., 2024; Borcherding et al., 2024). Several questions remain unaddressed, however, including the early cellular kinetics of LN activation, and its coordination across LNs located at different anatomical sites. It is not understood how this process is influenced by an adjuvant, such as MF59C.1, used in adjuvanted quadrivalent influenza vaccine (aQIV). In the event of an avian influenza pandemic, an H5N1 influenza vaccine including the MF59C.1 adjuvant could be deployed, but how this formulation induces immunity in young adults is not well studied, as it is not used in this population for seasonal immunisation.

Another fundamental gap in human LN data is the lack of representation from individuals of diverse, non-white European ethnicities, including those of African and Asian ancestry. Variation in inflammatory and immune responses in individuals with different ethnic backgrounds has been documented (Coussens et al., 2013; Mostafavi et al., 2016), with likely implications for the susceptibility and severity of response to pathogens. With respect to globally prevalent seasonal infections, such as influenza, respiratory mortality associated with the virus is highest in sub-Saharan Africa (2.8-16.5 per 100,000 individuals) and South-East Asia (3.5-9.2 per 100,000 individuals) (Iuliano et al., 2018). This may be attributed to geographic, socioeconomic and/or genetic factors, but notably, the risk of influenza infection and of severe outcomes following infection is also higher in individuals of Black, South Asian or mixed ethnicity residing in England (Zhao et al., 2015; Davidson et al., 2021) and in the United States of America (USA) (Chandrasekhar et al., 2017). Similarly, non-white ethnicities were disproportionately represented amongst working-age adults with severe sequelae from the COVID-19 pandemic in the United Kingdom (UK) and USA (Sze et al., 2020; Mathur et al., 2021). Research addressing the immunological factors that underpin these differences - such as population-dependent human leukocyte antigen (HLA) diversity, variation in immune cell activation in LNs, and the orchestration of responses across the lymphatic network - is required to inform the design of vaccines and immunotherapeutics to help overcome health vulnerability and to boost pandemic preparedness.

To address the paucity of LN data from individuals of non-white ethnicity, and from multiple LN sites sampled simultaneously, we have profiled LNs before and after vaccination with the 2022-23 seasonal quadrivalent influenza vaccine adjuvanted with MF59C.1 (aQIV) in a cohort of young adults of African or Asian ancestry recruited from the West London area in the UK as part of the ‘Lymph nodE single-cell Genomics AnCestrY’ (LEGACY) Network. FNA was performed to obtain cells from both the axillary draining and non-draining LNs (dLNs and ndLNs, respectively) for analysis by single cell multiomics. Our study thus provides an important resource with ethnic representation of lymph node cells at the genetic, transcriptomic and protein level, and shows the early anatomical, temporal and cellular dynamics of human axillary LNs after intramuscular immunisation in ancestrally diverse young adults.

## Results and Discussion

### Rarely studied HLA types amongst ancestrally diverse participants responding to aQIV

Adult volunteers with self-declared African or Asian ancestry (n=13) were enrolled and completed visits during the northern hemisphere winter flu season of 2022/23 **(Fig. 1A).** One additional participant was enrolled but did not complete the study due to contracting COVID-19. To collect steady state lymph node cells (LNC) ultrasound (US)-guided fine-needle aspiration (FNA) of axillary lymph nodes in both the left and right axillae was performed with paired collection of peripheral blood, prior to or at the start of the 2022/23 winter season. Participants received the adjuvanted quadrivalent influenza vaccine (aQIV) by intramuscular injection into the deltoid muscle of the arm. Arm selection for injection (left or right) was by participant choice and was recorded by the study team. Five (+/-2) days later US-guided FNA of draining lymph nodes (dLN) and non-draining lymph nodes (ndLN) was repeated with a final collection of serum at 28 days after immunisation **(Fig. 1B).**

**Figure 1.**
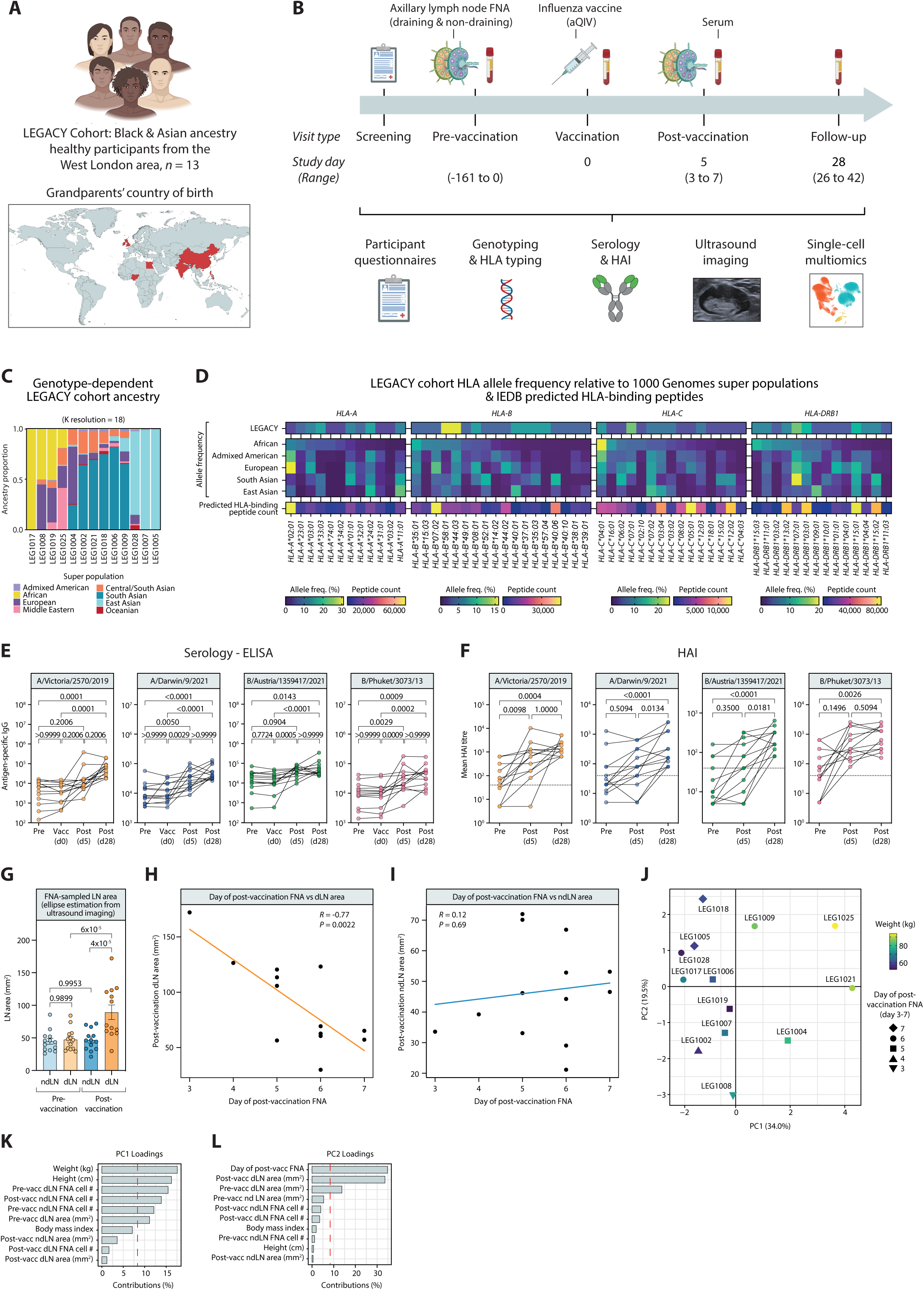
LEGACY study design, cohort ancestry and serological and anatomical response. **(A)** West London cohort of healthy participants who self-identified as having Black and Asian ancestry (n = 13). Global map indicates the country of birth of the participants’ grandparents. **(B)** Timeline of the study where multi-modal data is collected at various time points before and after influenza vaccination (aQIV: adjuvanted quadrivalent influenza vaccine). **(C)** Cohort ancestry as determined by genotype-dependent admixture analysis. Stacked bars represent the admixture composition of each participant at K=18. **(D)** Cohort HLA allele frequency relative to 1000 Genomes super populations and the count of peptides predicted to bind to these alleles based on the Immune Epitope Database (IEDB). **(E)** Detection of IgG antibodies against influenza strains used in the 2022/23 influenza vaccine by ELISA assay at various timepoints (pre: pre-vaccination, vacc: day of vaccination, post (d5): median five days post-vaccination, post (d28): median 28 days post-vaccination). Differences in IgG levels by timepoint were assessed by the Friedman test. **(F)** Haemagglutination inhibition (HAI) assay to identify functional antibody responses to the viruses used in the 2022/23 influenza vaccine. The dotted line represents the HAI seroconversion threshold (HAI≥40). Differences by timepoint were assessed by the Friedman test. **(G)** Area of FNA-sampled draining and non-draining lymph nodes (dLN and ndLN, respectively) before and after vaccination. Error bars represent the standard error of an ellipse. One-way ANOVA compared effect of sample location and time point with LN size. There was a significant area difference between at least two groups (F = 12.914, *P* = 2.61 x 10^-6^). A Tukey HSD post-hoc test for multiple comparisons found that the mean of post-vaccination dLN area was significantly different compared to the other sample location and time points (*P* < 0.0001). **(H,I)** Linear correlation of **(H)** dLN and **(I)** ndLN estimated area (mm^2^) with the number of days post-vaccination. *R* is the correlation coefficient, *P* is the *P*-value. **(J-L)** Principal component analysis (PCA) was conducted on pre- and post-vaccination (pre-vacc and post-vacc, respectively) LN cell counts, LN area (as estimated from ultrasound images), and donor characteristics such as weight (kg), height (cm), and BMI. **(J)** PCA coloured by donor weight (kg). Symbols indicate the day of post-vaccination FNA for each participant. Principal component (PC)1 and PC2 percentages indicate the variation captured by each component. **(K,L)** PC1 and PC2 loadings. Dashed red threshold line represents the theoretical value if all variable contributions were equal.

The study was sponsored by Imperial College London and approved by the Research Ethics Committee (REC ref: 22/LO/0343). Participants provided written, informed consent and were monitored for safety throughout; there were no study vaccine related serious adverse events. Further details are available in the methods, protocol (Pollock & Dendrou, 2023) and the study registration (ISRCTN13657999).

Participants were 5 men and 8 women, aged 34 years (+/-4.8 years). Participants reported a wide range of self-declared ethnicity including Middle Eastern, Asian, Black African, West African and mixed ethnicities. The country of birth was most frequently United Kingdom/England, with others reported as France, China, India and Trinidad and Tobago. All participants had been resident in the UK for at least 5 years before enrolling into the study. The parents’ and grandparents’ country of birth collectively included 11 unique countries **(Table S1)**. Consistent with this, the ancestral diversity of the participants was evident through genome-wide genotyping and admixture analyses, demonstrating representation from across eight different ancestral super populations **(Fig. 1C)**.

Participants also showed human leukocyte antigen (HLA) genotype diversity, with several rarely studied HLA genotypes being present in the cohort, as compared to the 1,000 Genomes Project super population cohorts (Abi-Rached et al., 2018). HLA class I and II alleles were highly polymorphic across the cohort; for example, for the *HLA-B* locus, 20 discrete alleles identified, of which 13 (65%) were rarely described alleles **(Fig. 1D; Table S2**). Given this HLA allele heterogeneity, the count of peptides predicted to bind to the different HLA alleles in the cohort, based on the Immune Epitope Database (Vita et al., 2019), varied widely across four orders of magnitude **(Fig. 1D)**. This demonstrates the challenge of predicting peptide binding for HLA alleles more prevalent in ancestrally diverse populations which are poorly represented in databases: computational prediction of HLA epitope binding is critical to machine learning (ML) approaches employed to aid rational vaccine design, but such ML models are biased as the majority of input data are derived from individuals of European ancestry (Bravi, 2024).

### Temporal regulation of binding and functional serological responses

Each aQIV 2022-2023 influenza vaccine administered to the LEGACY study participants contained four 15 microgram doses of the homotrimeric glycoprotein haemagglutinin (HA), raised from A/Victoria/2570/2019 IVR-215, A/Darwin/6/2021 IVR-227, B/Austria/1359417/2021 BVR-26 and B/Phuket/3073/2013 BVR-1B. Participant responses to the vaccine were measured by means of binding Ig ELISA, Luminex and haemagglutination inhibition (HAI) assays, using participant sera obtained before, upon and after vaccination **(Fig.1E,F; Fig. S1)**. At both post-vaccination time points there was a significant increase in antigen-specific IgG to all four-vaccine virus strain HAs **(Fig. 1E)**. The IgG response was largely IgG1, whilst IgG3 also contributed towards A/Darwin and A/Victoria responses at the earlier time point, 5 days post vaccination **(Fig. S1)**. There was a rapid IgM response at day 5 which waned by day 28 and rapid IgA response which was sustained against the vaccine antigens **(Fig. S1)**. Although not included in the vaccine, there was also an increase in IgG towards both avian HA subtypes H5N1 and H5N8 at 28 days post vaccination **(Fig. S1)**. Where there was no pre-existing protection, participants seroconverted (HAI ≥40) against all four vaccine antigens **(Fig. 1F)**.

### LN cellular composition is both temporally and anatomically regulated

During FNA visits, B-mode US images were taken of dLNs and ndLNs and measurements taken of the long and short axis. Prior to vaccination during steady state, dLN and ndLN areas averaged 47.4 and 44.8 mm^2^, respectively and did not differ significantly. Post-vaccination, there was a significant, approximately 2-fold increase in dLN area. However, there was no significant detectable change in the ndLN size (**Fig. 1G**). The post-vaccination dLN size was significantly negatively correlated with the number of days after vaccination, but this was not the case for ndLN size (**Fig. 1H,I**).

Further to investigating LN size changes, we also assessed the yield of LNCs from the US-guided FNA. Yields were variable, but in the same range as previous studies (Turner et al., 2020; Day et al., 2022). To determine which factors were contributing to this variation, a principal component analysis (PCA) was undertaken. Donor quantitative factors such as height and age, the total cell count after sample processing, the study time points, alongside LN size were considered. The number of days post vaccination contributed significantly to the cell yield. Donor height and weight were also strong contributors to LN cell yield, especially at steady state prior to vaccination **(Fig. 1J-L)**.

### A longitudinal multi-modal scRNA-seq vaccination atlas of axillary dLNs and ndLNs

Single-cell analysis of 45 LN FNA samples was undertaken from the 13 participants who completed the study protocol, resulting in 191,266 high-quality transcriptomes alongside cell-surface protein and T-cell repertoire data. Three main cell compartments were identified, T/NK/IL, B/plasma, and non-lymphoid cells, and were subclustered into 42 different cell states: 20 T and NK cell states (n=99,212 cells), 10 B cell and plasma cell states (n=86,137 cells) and 12 non-lymphoid (myeloid and stromal) cell states (n=5,917 cells) **(Fig. 2A-D)**. Amongst these, cell states important for initiating and sustaining the serological vaccine reaction within LNs were readily identified **(Fig. 2E-G)**. The identity of these cell states, which included CD4^+^ T follicular helper (Tfh) cells, germinal centre (GC) light zone (LZ) B and cycling B cells, plasma cells and plasmacytoid dendritic cells (pDCs), were confirmed by cell-surface protein marker expression based on antibody-derived tags **(Fig. 2E-G)**. For example, Tfh cells expressed marker transcripts *CD3E*, *CD4*, *CD28*, *CD69*, *CXCR5*, *PDCD1*, *BCL6* and *CD44*. The same cell state also expressed the protein markers CD3, CD4, CD45RO, PD-1, ICOS, CXCR3, CXCR5, B and T lymphocyte attenuator (BTLA) and CD57 **(Fig. 2E)**. Other notable non-naïve lymphocyte cell types detected included CD8^+^ *GZMK*^+^ T cells, CD8^+^ *GZMB*^+^ T cells and mucosa associated invariant T (MAIT) cells. Non-lymphoid cells were less abundant but highly diverse, particularly amongst dendritic cells which included six different identifiable cell states **(Fig. 2D,G)**.

**Figure 2.**
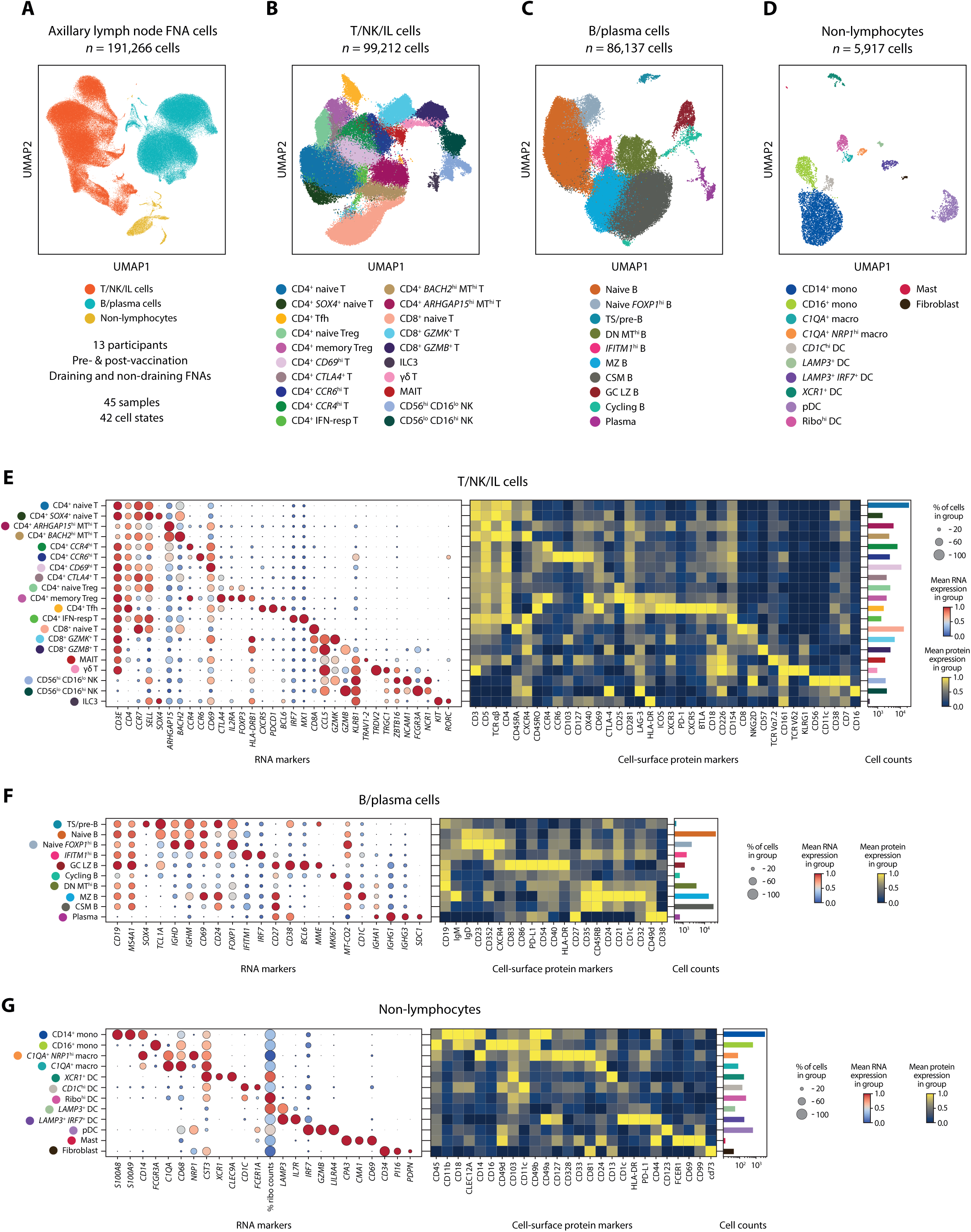
Characterisation of LN FNA cell populations and cell states identified using single-cell transcriptomics and surface protein expression. **(A)** Uniform Manifold Approximation and Projection (UMAP) representation of the three broad cell compartments identified in LN FNA samples pre- and post-vaccination. **(B-D)** Higher resolution annotation of T/NK/IL and B/plasma cell and non-lymphocyte compartments. CSM, class-switched memory B cell; DC, dendritic cell; DN, double negative B cell; FNA, fine-needle aspirate; GC LZ, germinal centre light zone B cell; hi, high; IFN-resp, interferon-responsive; IL, innate lymphoid; lo, low; macro, macrophage; MAIT, mucosal-associated invariant T; mono, monocyte; MT, mitochondrial genes; MZ, marginal zone B cell; NK, natural killer; pDC, plasmacytoid dendritic cell; Ribo, ribosomal; Tfh, T follicular helper cell; Treg, T regulatory cell; TS, transitional B cell. **(E-F)** Key RNA marker genes (left), cell-surface protein expression (middle), and cell counts (right) of each cell state per cell compartment.

### Influenza vaccination induces anatomical and temporal-specific differential cellular abundance

Adjuvants generate a robust immune response in a non-antigen specific manner; the MF59C.1 adjuvant in the vaccine received by the LEGACY study participants is known to support early generation of high-affinity influenza-specific antibodies and circulating Tfh in older adults (Li et al., 2021). Thus, MF59C.1 potentially affects the dLNs and ndLNs similarly, depending on its pharmacokinetics. Furthermore, cellular infiltration of non-resident memory B cells may be required to support the LN recall response, as shown in mice (Mesin et al., 2020). To determine whether an adjuvanted influenza vaccine induces cellular immunity across the lymphatic network, or regionally, we investigated the differential abundance changes in the dLNs and ndLNs before and after vaccination **(Fig. 3A)**. To utilise the multiple sample-sites per donor as a means of within-donor normalisation, mixed-effects association testing for single-cells (MASC) (Fonseka et al., 2018) was used to test differential abundance between sample sites and time points for each cell state directly, while accounting for weight and number of days post-vaccination as fixed-effect covariates. Overall, the differential abundance changes in the dLN after vaccination were dramatically different to those in the ndLN **(Fig. 3A)**. These results were mirrored in the direct post-vaccination dLN and ndLN comparison. CD4^+^ Tfh cells were significantly higher post-vaccination on the dLN side, but not the ndLN side compared to pre-vaccination. Like our previous findings, naïve B cells, germinal centre and cycling B cells, and CD4^+^ CD69^+^ T cells increased in abundance in dLNs (Day et al., 2022). Changes in T-cell abundance in ndLNs were amongst cells not typically associated with antigen-specific responses, including CD4^+^ IFN-resp T, MAIT and NK cells. In both dLNs and nd LNs, plasma cells, cycling B cells and pDCs were relatively increased, as were CD4^+^ memory Treg cells, with relative decreases in CD8^+^ GZMB^+^ T-cell abundance. This suggested a second, less prominent but measurable global effect of aQIV immunisation on the lymphatic system, that could potentially be attributed to the adjuvant, with highly abundant vaccine-responsive cell types, e.g., CD4^+^ Tfh, diluting this effect in the dLN. The median proportion of each cell state at each sample site and time point further illustrates these findings **(Fig. 3A)**.

**Figure 3.**
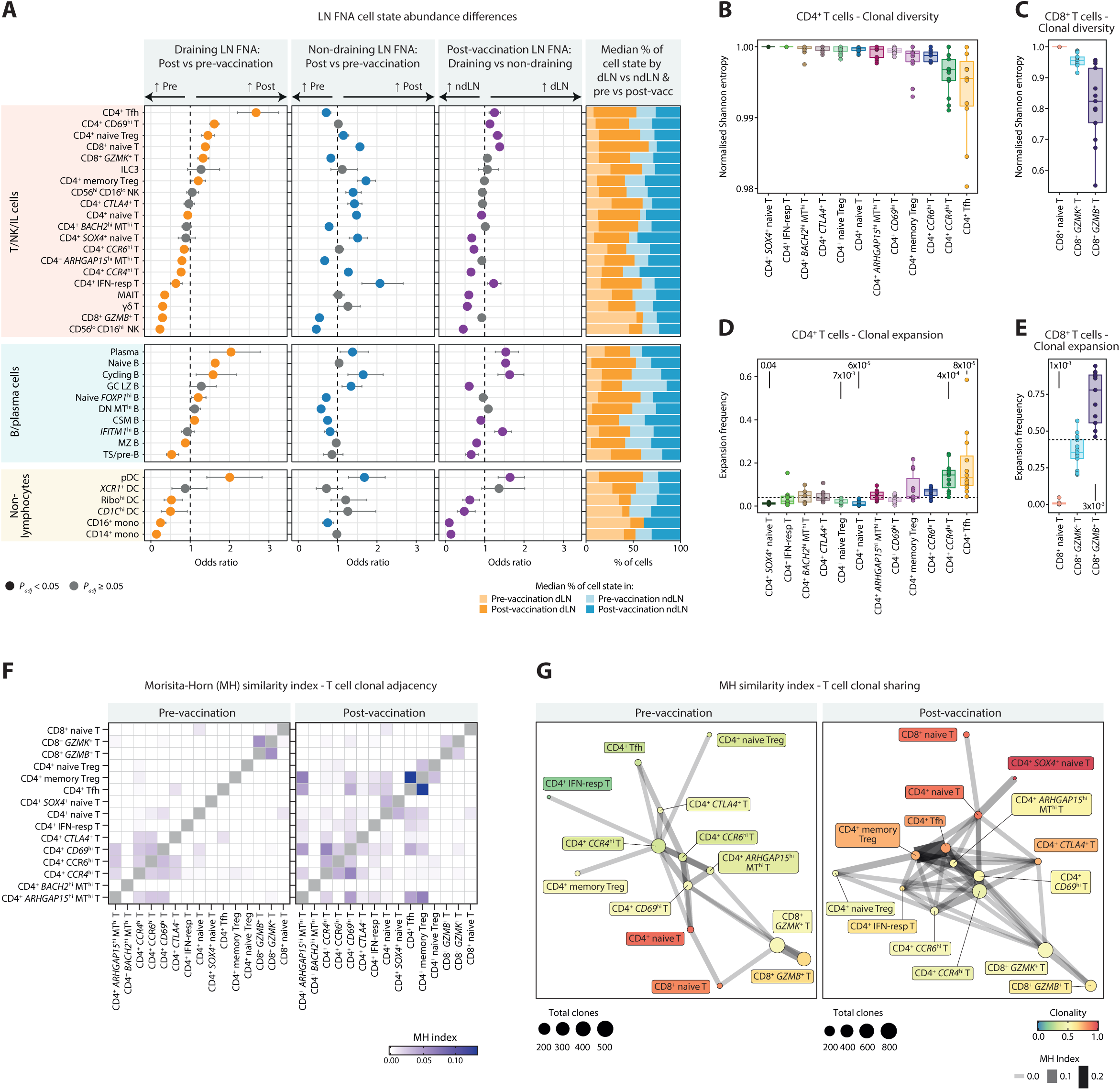
Influenza vaccine stimulates differential abundance and T-cell clonal dynamic changes across cell states, sample sites and time points. A total of 80,427 cells with both transcriptomic and TCR sequences were observed across all subsets, sample sites, and donors including 2,057 clones (groups of two or more cells sharing TCRs). **(A)** MASC used to test abundance differences across cell compartments after influenza vaccination (see Methods for more details). Comparison of pre-vaccination and post-vaccination samples from draining LNs (dLNs; orange; pre-vaccination: n = 10, post-vaccination: n = 12) and non-draining LNs (ndLNs; blue; pre-vaccination: n = 11, post-vaccination: n =12) as well as a post-vaccination comparison of dLNs and ndLNs (purple) represented as odds ratios (OR) are shown. Non-grey dots indicate significant differences (*P_adj_* < 0.05) after Benjamini-Hochberg correction. Error bars show the 95% confidence interval. Median proportion of each cell state by sample site and time point is also represented in a stacked bar plot. **(B-C)** Normalised Shannon entropy index for each cell state in CD4^+^ or CD8^+^ T-cell compartment. Clonotypes included have one alpha and one beta chain. Each dot represents a donor. **(D-E)** Proportion of cells in a clone (group with two or more cells) in each cell state in CD4^+^ or CD8^+^ T-cell compartment. Clonotypes included have one alpha and one beta chain. Dashed line indicates the mean expansion frequency in that compartment. Wilcox test compared the expansion frequency of each cell type to the mean of the CD4^+^ or CD8^+^ T-cell compartment expansion frequency. **(F)** Clonal dynamics between cell types pre-vaccination and post-vaccination as indicated by Morisita-Horn (MH) similarity index. **(G)** Shared clonal networks of pre-vaccination and post-vaccination T-cell repertoires after filtering where edge weights are defined by MH similarity index, node colour denotes clonality, and node size indicates number of clones. CSM, class-switched memory B cell; DC, dendritic cell; DN, double negative B cell; FNA, fine-needle aspirate; GC LZ, germinal centre light zone B cell; hi, high; IFN-resp, interferon-responsive; lo, low; macro, macrophage; MAIT, mucosal-associated invariant T; mono, monocyte; MT, mitochondrial genes; MZ, marginal zone B cell; NK, natural killer; pDC, plasmacytoid dendritic cell; Ribo, ribosomal; Tfh, T follicular helper cell; Treg, T regulatory cell; TS, transitional B cell.

### Vaccination-induced early changes in LN T-cell repertoire

We next investigated the dynamic changes in the T cell repertoire upon vaccination to further elucidate the relationships between T cells. Using Immcantation, IgBlast and scRepertoire framework for V(D)J gene assignment and productive alpha-beta paired clonotype filtering, clones were identified using VDJC genes and the CDR3 nucleotides on both alpha and beta chains. Normalised Shannon entropy index revealed a relative drop in the diversity of CD4^+^ Tfh, CD4^+^ *CCR4*^hi^ T cells, and CD8^+^ *GZMB*^+^ T cells **(Fig. 3B,C)**. Similarly, the expansion frequency was also significantly increased in the same cell states compared to the median expansion frequency of CD4^+^ and CD8^+^ T cells respectively **(Fig. 3D,E)**. Collectively, this suggests that the increase of CD4^+^ Tfh cells in LNs is not only a reflection of increased recruitment from the circulation, but also expansion and differentiation. Morisita-Horn similarity index was used to compare the T-cell immune repertoire before and after vaccination among different T-cell states. Post-vaccination clonal dynamics were found to be more active than pre-vaccination, especially in the CD4^+^ T-cell compartment **(Fig. 3F,G)**. CD4^+^ *CCR4*^hi^ T cells, which showed small but significant changes in abundance bilaterally had heavily expanded TCRs and after vaccination this cell type remained a key “hub”, showing the most clonal connections to other cells and an increased clonal link to CD4^+^ *CD69*^hi^ T cells post vaccination **(Fig 3G)**. The role of CD4^+^ *CCR4*^hi^ T cells in vaccine responses has not previously been identified in humans *in vivo*; CCR4 affects T-cell LN homing and thymocyte recruitment to the site of immunisation in mice (Palchevskiy et al., 2019; Yamamoto et al., 2018) and CCR4 antagonists target cells with adjuvant activity *in silico* (Bayry et al., 2008).

### Influenza vaccination stimulates gene expression hubs associated with dLNs but not ndLNs

To further investigate the heterogenous immune response after influenza vaccination, we utilised consensus non-negative matrix factorisation (cNMF) to identify gene expression programmes (GEPs) within cell types (Kotliar et al., 2019). Considering a cell’s gene expression can reflect its cell type concurrently alongside its metabolic and functional state, any individual cell can comprise multiple GEPs **(Fig. 4A)**. After identifying the GEPs of each broad compartment (CD4^+^ T cells, CD8^+^ T/NK/IL cells, B/plasma cells, and non-lymphocytes), we assessed the contribution of each GEP to each cell type **(Fig. 4B)** and top genes/pathways to each GEP **(Fig. 4C)**. For instance, CD4^+^ Tfh cells comprised of multiple GEPs including GEPs pCD4T_9 and pCD4T_14 that identified the cell type with genes such as *PDCD1* and *CXCR5*, as well as shared GEPs pCD4T_2 and pCD4T_3 that mapped to many other CD4^+^ T cell states and included genes such as *IL7R*, *CD96*, and ribosomal genes.

**Figure 4.**
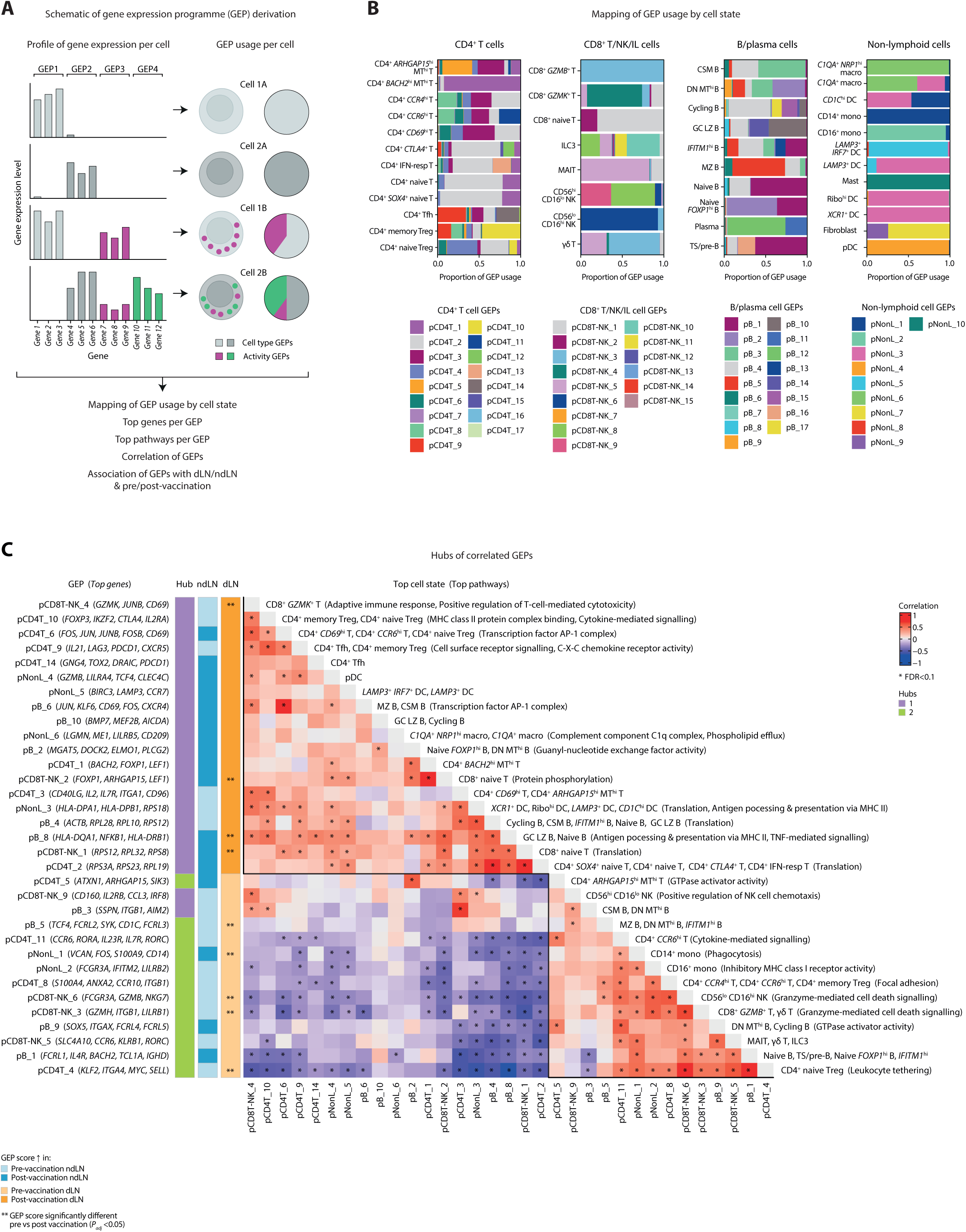
Differential gene expression hubs associated with influenza vaccination. **(A)** Schematic of cNMF analysis identifying gene expression programmes (GEPs), their characterisation, correlation between significant GEPs, and association with sample site and time points. **(B)** Mapping of GEP usage by cell state across CD4^+^ T cell, CD8^+^ T/NK/IL cell, B/plasma cell, and non-lymphoid cell (Non-L) compartments. **(C)** Correlations of significant GEPs with annotations of top genes, pathways and cell types. * indicates significantly correlated GEP pairs (FDR < 0.1). Solid lines demarcate hubs of highly correlated GEPs. Hub annotation indicates two clusters from hierarchical clustering. dLN and ndLN annotations denote which GEPs have greater expression at each sample site and time point. ** indicates the GEPs that have significantly different expression pre- vs post-vaccination (*P_adj_* < 0.05). CSM, class-switched memory B cell; DC, dendritic cell; DN, double negative B cell; FNA, fine-needle aspirate; GC LZ, germinal centre light zone B cell; hi, high; IFN-resp, interferon-responsive; IL, innate lymphoid; lo, low; macro, macrophage; MAIT, mucosal-associated invariant T; mono, monocyte; MT, mitochondrial genes; MZ, marginal zone B cell; NK, natural killer; pDC, plasmacytoid dendritic cell; Ribo, ribosomal; Tfh, T follicular helper cell; Treg, T regulatory cell; TS, transitional B cell.

Significant GEPs were then further correlated with each other to identify hubs of related biological processes. We derived two hubs using hierarchical clustering **(Fig. 4C)**. We assessed how the individual GEPs changed pre- and post-vaccination in the dLNs and ndLNs. GEPs associated with the hub 1 had a higher median expression post-vaccination on the dLN side but not the ndLN side. Post-vaccination GEPs in hub 1 included genes associated with CD4^+^ Tfh and CD4^+^ memory T reg, pDC and cycling B cells. On the other hand, GEPs associated with hub 2 had increased median expression pre-vaccination on the dLN side, but not the ndLN side. In summary, the early vaccine response gene signature to adjuvanted QIV can be detected in the dLN. This vaccine-specific gene signature mapped directly to the LN response to aQIV in healthy young adults with diverse ancestry, was associated with robust vaccine immunogenicity and could serve as benchmark for comparison of vaccine-induced immunity with other platforms or in elderly or immunosuppressed populations.

This study demonstrated that the early (within one week) lymphatic response to intramuscular immunisation with aQIV was tightly temporally, genetically and regionally regulated. This response pattern was broadly consistent across young adults receiving MF59C.1 aQIV and was not strongly influenced by ancestral diversity, despite vast differences in genotype and HLA type. Using US to serially track responses, immunisation into the deltoid muscle was associated with a fast onset of regional swelling in the dLNs only, with an indication from later stage data, that this swelling might resolve equally rapidly. This early swelling was accompanied by an influx of adaptive immune cells involved including CD4^+^ Tfh and B cells, with a drop in clonal diversity indicating T-cell clonal expansion is an active process within the dLN. This finding builds on recent work on later-stage cellular events in the dLN with non-adjuvanted influenza vaccine (Schattgen et al., 2024), and we demonstrate that dLN swelling can be directly linked to cellular infiltration and expansion, consisting of multiple vaccine-responsive cell types dominated by CD4^+^ Tfh in the first few days post vaccination. Cellular dynamics of the ndLN have not been previously studied by FNA and single-cell multiomics, but our data indicate that the coordinated swelling and adaptive immune cell increase does not occur in this time frame, although some cellular changes are observed after vaccination.

Here we have shown the cellular and genetic determinants of the regionally and temporarily defined early lymphatic response to intramuscular immunisation with an adjuvanted quadrivalent influenza vaccine, in a highly ancestrally diverse cohort of young adults. This dataset serves as a publicly available resource to address inequity in reference material underpinning basic and translational discoveries in human immunology.

## Materials and methods

### Clinical study

#### Registration and design

The full version of the approved clinical study protocol, version 2.1 is available (Pollock & Dendrou, 2023). The study is registered ISRCTN13657999 (https://doi.org/10.1186/ISRCTN13657999). The study was approved by the London - Central Research Ethics Committee. The Sponsor was Imperial College London. This was a single-site interventional non-randomised open label experimental medicine clinical research study in thirty volunteers in total over two influenza seasons (2022-2023 and 2023-2024). The study was run at the National Institute for Health and Care Research (NIHR) Imperial Clinical Research Facility, West London, UK. We report here data from the first season cohort, who completed all study visits in 2023-2023 (n=13).

#### Eligibility and enrolment

Participants were eligible to volunteer for the study if: (i) they were healthy adults aged ≥18 years and ≤55 years on the day of screening, (ii) were willing and able to provide written informed consent, identified as having African or Asian ancestry, (iii) were usually resident in the UK for ≥5 years prior to screening, (iv) were not pregnant on the day of screening and were willing to use a highly effective form of contraception until 12 weeks after the study immunisation (if a person of childbearing potential), (v) were willing to avoid all other vaccines within 4 weeks either side of the study injection and fine needle aspiration, (vi) were willing and able to comply with the visit schedule and provide samples, and (vii) were willing to grant authorised persons access to their trial related medical record and GP records either directly or indirectly. Participants were given written information about the study and the opportunity to ask questions. Those wishing to take part gave written informed consent and underwent screening procedures including a general health check, medical history assessment and blood tests to confirm eligibility. Participants who were pregnant or lactating, had a significant medical history, a body mass index of ≥30, a history of allergy to drugs/vaccines, anaphylaxis or angioedema, had recently (within 18 weeks) taken immunosuppressive agents, or were taking part in another trial were not eligible. Participants underwent screening tests for blood borne viruses and were ineligible if they had detectable antibody to HIV, hepatitis C or hepatitis B.

#### Safety

There were no safety objectives of this experimental medicine study. To ensure the safety and well-being of participants safety assessments were conducted as follows. During the study, participants were asked about concomitant medications at every visit and about COVID-19 symptoms at every in person visit. People of childbearing potential had a urinary pregnancy test at screening and on the day of immunisation and were asked to use an effective form of contraception during the study if sexually active. A symptom directed physical examination was performed where necessary. Adverse events were recorded in the first 5 days following LN tissue sampling and in the 28 days following influenza vaccination for the study. Serious adverse events were collected throughout the study. Side effects of the vaccination that were suspected adverse reactions were reported to the MHRA through the yellow card scheme by the investigator.

#### Study immunisation

The study immunisation, adjuvanted quadrivalent influenza vaccine (aQIV) (CSL Seqirus), was administered as an intramuscular injection into the non-dominant arm (as requested by study participants) in the deltoid muscle. One 0.5 mL dose of aQIV contained 15 µg of haemagglutinin from two A and two B strains of influenza propagated in hens’ eggs and adjuvanted with MF59C.1. This contained per 0.5mL dose, squalene (9.75 mg), polysorbate 80 (1.175 mg), sorbitan trioleate (1.175 mg), sodium citrate (0.66 mg) and citric acid (0.04 mg). The 2022/2023 UK recommendation for influenza vaccine study composition was: A/Victoria/2570/2019 (H1N1)pdm09-like strain, A/Darwin/9/2021 (H3N2)-like strain, B/Austria/1359417/2021-like strain, and B/Phuket/3073/2013-like strain. The aQIV clinical study vaccine contained: A/Victoria/2570/2019 IVR-215, A/Darwin/6/2021 IVR-227, B/Austria/1359417/2021 BVR-26, and B/Phuket/3073/2013 BVR-1B.

#### Ultrasound-guided fine-needle aspiration sampling procedure

The US-guided FNA was taken by a qualified medical practitioner according to our established SOP. Participants were ineligible to undergo the procedure if they were taking blood thinning medication prior to aspiration, if there were signs of local infection, or if there was pain or swelling at the site of sampling. The FNA was conducted using standard aseptic technique under US guidance using a Toshiba Aplio i700 US machine. Following infiltration of local anaesthetic, a sterile needle and syringe were inserted into the axillary lymph node under US guidance using 3-5 passes. Aspirated cells were placed directly into R10 solution for transport to the laboratory in a sealed air-sea container with cooled gel packaging.

#### Sample processing

Samples were processed immediately on receipt, as previously described for viable cryopreservation (Day et al., 2022; Pollock and Dendrou, 2023).

Lymph node (LN) FNA samples were processed as previously published (Day et al., 2022). In brief, LN FNAs were collected in R10 media and transported at 4°C to the laboratories. R10 media contains RPMI 1640 with 25 mM HEPES (Cat No: R5886, Sigma), 10% heat-inactivated fetal bovine serum (HI-FBS) (Cat No: F4135, Sigma), 1% Pen/Strep (Penicillin-Streptomycin 10,000 Units/mL (Cat No: 15140-122, Gibco), and 2 mM of L-Glutamine (Cat No: 25030-024, Gibco). Samples were washed in RPMI with 5% human AB serum, incubated in ACK lysis buffer (Gibco, Cat: A10492-01) at room temperature for 5-10 minutes, and then washed twice with cell wash buffer (PBS + 2% human AB serum + 2 mM EDTA). Cells were then cryopreserved in CS10 in aliquots of 500,000 cells in 100 µL.

#### Ultrasound analysis

Images of sampled LNs, typically six per axillae, were taken at the FNA visits before and after immunisation. Analysis of images was undertaken by two medical practitioners confirming the clearest image of the sampled node which was then measured in two axes in the cross-sectional plane.

#### Genotyping, HLA typing and HLA peptide-binding estimation

DNA was isolated from whole blood using the QIAamp DNA Blood Midi Kit according to manufacturer’s instructions. Genotyping was performed using the Infinium Global Screening Array-24 v3.0 (Illumina) by the Genome Centre, Queen Mary University of London according to manufacturer’s recommendations. This dataset contained 730,000 polymorphic variants. HLA typing, specifically HLA class I HLA-A/B/C (exons 2 and 3) and class II DRB1/DQB1 (exon 2 only) up to 4 digits, was performed by the sequencing/typing facility at the MRC Weatherall Institute of Molecular Medicine, University of Oxford. LEGACY cohort HLA allele frequency was compared to the 1000 Genomes Project HLA data (Abi-Rached et al., 2018) derived from five ancestral super populations (African, *n*=712; Admixed American, *n*=373; European, *n*=529; South Asian, *n*=543; and East Asian, *n*=536 individuals). The count of peptides predicted to bind to different HLA-A/B/C and HLA-DRB1 alleles was obtained and plotted from estimates provided by the Immune Epitope Database (Vita et al., 2019).

#### Single-cell experiments

LN FNA samples, pre-stained with TotalSeq-C Human Universal Cocktail V1.0 (BioLegend) when cell numbers allowed (as per the manufacturers’ guidance), were loaded with a maximum of 20,000 cells onto 10x Genomics Chromium X controller with standard kits. Libraries (5’ gene expression, V(D)J, and ADT) were prepared using manufacturers’ guidelines and sequenced using NovaSeq 6000 PE150 for LN FNA and NovaSeqX Plus PE150 (NovoGene) for PBMCs.

#### Serological analysis

Antibodies raised in serum by the influenza vaccine were measured by means of a binding antibody ELISA and Luminex assays. Friedman tests were used to assess statistical significance in antibody levels by timepoint, with a significance threshold of *P*-value < 0.05.

#### Haemagglutination Inhibition

Serum samples were analysed by Haemagglutination Inhibition (HAI) with the viruses used in the 2022/23 influenza vaccine: A/Victoria/2570/2019 (H1N1)pdm09-like strain, A/Darwin/9/2021 (H3N2)-like strain, B/Austria/1359417/2021-like strain, B/Phuket/3073/2013-like strain. The principle of the HI test is based on the ability of specific anti-influenza antibodies to inhibit hemagglutination of red blood cells (RBCs) by influenza virus HA. Serum was pre-treated with neuraminidase followed by heat inactivation for 1h at 56°C. Samples were titrated in an 8-step two-fold dilution series, incubated with the HA antigen suspension for 1 hour at room temperature, and 25 µL of the 0.5% RBC suspension (turkey blood for the H1N1 and B strains; guinea pig blood for the H3N2 strain) added. The reaction was left for up to one hour before reading.

Titre duplicates for each serum-virus combination achieved less than 2-fold variance or were repeated. The standard dilution series covered titres 10-1280 with samples which did not reach an endpoint in this initial titration undergoing endpoint titration; titres which were considered to indicate sero-protection were HAI titres ≥40. Friedman tests were used to assess statistical significance by timepoint, with a significance threshold of *P*-value < 0.05.

### Computational

#### Ultrasound images, correlation and principal component analysis (PCA)

Area of lymph nodes were estimated as an ellipse from the width and length measurements of the lymph nodes. Standard error for the area was calculated as

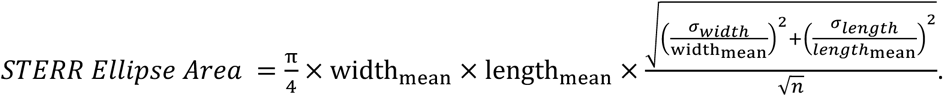

Outliers, determined by 1.5*interquartile range beyond first and third quantile, were removed prior to linear correlation analysis of estimated lymph node area and number of days post-vaccination. Principal component analysis was performed using donor characteristics, sampling cell counts, and estimated lymph node area. Prior to performing PCA, missing data was imputed using imputePCA (missMDA package) with number of components used equal to half the number of variables used in the PCA. PCA results were visualised using factoextra package.

#### Genotyping quality control and admixture

The input array data consisted of 13 samples and 730,000 variants prior to filtering and QC. Filtering and QC steps included: 1) retaining only autosomal variants, 2) removing loci where 99.9% of genotypes are missing, 3) removing SNPs with minor allele frequency of 1%, and 4) excluding variants with one or more multi-character allele code. The final post-QC array data consisted of 330,000 variants. The reference dataset of merged genotypes combined samples from 1000 Genomes and HGDP. Intersecting variants between our data and the reference data for downstream analysis was determined by the following: 1) exclude non-unique SNPs, 2) exclude A-T or G-C SNPs, and 3) aligned position mismatches and allele flips. The program ADMIXTURE (v.1.3.0) was used to estimate per-individual ancestry populations in a panel of 3433 reference individuals representing African, European, East Asian, and American ancestries. The optimum number of ancestry populations (K) was chosen based on fivefold cross-validation for each K in the set of 5-30. Model of K = 18 had the lowest cross-validation error. The population allele frequencies estimated from the analysis of reference samples were fixed as parameters so that the LEGACY samples could be projected into the admixture model to obtain ancestry proportion estimates.

#### Single-cell pre-processing, quality control to clustering and annotation

From the Chromium single-cell RNA sequencing outputs, Cell Ranger v7.0 was used for transcript alignment to GRCh38 version 2020-A and generation of feature-barcode matrices for downstream analysis. Panpipes was used for quality control, batch correction, dimension reduction, and clustering (Curion et al., 2024). High-quality single cells were identified by removing doublets determined using Scrublet (Wolock et al., 2019), cells expressing fewer than 300 genes, cells with mitochondrial gene count percentage greater than 25%, cells with haemoglobin gene count percentage greater than 40%. Genes detected in fewer than 3 cells were also removed.

UMI counts were normalised using the shifted logarithm technique found in scanpy’s pp.normalized_total function. Top 3000 highly variable genes, excluding *IGHV* and *TCR* genes, were selected using VST from Seurat v3 for downstream dimension reduction. Harmony was used for batch correction between experimental batches with 30 ncps. The results were then visualised in a lower dimension using UMAP. Leiden clustering helped identify three broad cell populations: T/NK/IL cells, B/plasma cells, and non-lymphocytes. These broad populations were then individually further clustered with re-calculated highly variable genes to generate finer annotations. Clusters with mixed gene signatures (ie. CD19 and CD3) were further clustered if possible, but otherwise removed if deemed high likelihood of being a doublet by the Scrublet score. The Wilcoxon rank-sum test was used to determine the top expressed genes in each cluster to aid with annotation.

#### Differential abundance analysis

MASC (mixed-effects modelling of associations of single-cells) (Fonseka et al., 2018) was used to identify differentially abundant cellular populations associated between conditions whilst accounting for participant’s weight and number days post-vaccination the second LN FNA sample was collected as fixed effects. Conditions compared include draining LN pre- vs post-vaccination, non-draining LN pre- vs post-vaccination, and post-vaccination draining vs non-draining LN.

#### Identification of gene expression programmes (GEPs) using consensus non-negative matrix factorisation (cNMF)

cNMF was used to simultaneously capture gene functional programs and activation states alongside cell identities (Kotliar et al., 2019). For each of the four broad cell types (B/plasma cells, CD4^+^ T cells, CD8^+^ T/NK cells, and non-lymphocytes), cNMF was iteratively run using 300 iterations and 3000 highly variable genes. The selection of k (number of components or gene expression programmes [GEPs]) was determined based on stability of the solution by the silhouette score, Frobenius reconstruction error, and biological relevance of the top weighted genes. Models of k = 17, 17, 15, and 10 were chosen for B/plasma cells, CD4^+^ T cells, CD8^+^ T/NK cells and non-lymphocytes, respectively. Outliers in consensus solutions were then filtered out based on histogram of components and their nearest neighbours using density thresholds of 0.15, 0.25, 0.25, and 0.21 for B/plasma cells, CD4^+^ T cells,, CD8^+^ T/NK cells and non-lymphocytes, respectively.

Conventionally, distributions are compared using single metrics like mean or variance. However, to directly compare distribution functions of program score expression, Wasserstein distance or Earth mover’s distance (EMD) was used to compute the difference between program score histograms between sample conditions (ie. visit type, vaccination side) with a bin size of 0.2. A large EMD score indicates a greater difference between the two sample distributions. To estimate the significance of the EMD scores for each program score per broad celltype, a permutation-based method was used to calculate a q-value. Specifically, using EMDomics package, the overall EMD score for each GEP is calculated by averaging all of the pairwise EMD scores and then labels randomly permuted 1000 times and the EMD score for each permutation is also calculated. The null distribution is determined by the median of the permuted scores for each GEP, and the FDR is determined for a range of thresholds. The thresholds that minimizes FDR is defined as the q-value. GEPs with significant q-values (q < 0.05) indicated that those gene programs have different distributions among the four sample conditions; thus, these GEPs were used in downstream hub analysis.

#### Identification of hubs and enrichment upon vaccination

Hubs with covarying GEPs were identified and compared between vaccination conditions. First, programme activity of each GEP per sample was summarised as a quantile mean (qmean) across five quantiles (0.25, 0.5, 0.75, 0.95, 0.99). A Pearson correlation co-efficient (R) was calculated for each GEP pairing across all samples. Fisher transformation was applied to the correlations, and the correlation mean was used as a test statistic. A null distribution was determined by permuting the sample identity 10,000 times whilst keeping the cell type constant, and R was tested against this null distribution. The correlation p-value was determined by counting how often the permuted R value differed from the true R value. The minimum count was scaled by two and designated the p-value statistic, and adjusted for multiple comparisons by Benjamini-Hochberg FDR of 10%. An adjusted R value was determined by calculating the difference of the mean true R values and the mean of permuted R values. Hierarchical clustering was used to cluster the GEPs. To determine if a GEP varied significantly before and after vaccination on the ipsilateral and contralateral side respectively, the qmean (as described above) of each GEP per donor per sample condition was compared using Wilcoxon rank-sum test and adjusted for multiple comparisons using Benjamini-Hochberg FDR of 10%.

#### Single-cell TCR analysis

The TCR repertoire analysis used the Immcantation and scRepertoire frameworks. V, D, and J genes were assigned using IgBlast. Non-productive sequences, cells with more than 3 alpha or beta chains, and those with gamma-delta constant regions were removed. Clones were identified using both VDJC gene and CDR3 nucleotide. Normalised Shannon entropy and expansion frequency (proportion of cells in a clone of two or more cells) were calculated using R package scRepertoire outputs.

To examine clonal overlaps of TCR repertoires between populations, we computed Morisita’s index using the R package divo for all pairwise T-cell clusters. For shared clonal network analysis and visualization, the R package igraph was used to construct weighted, undirected T-cell networks, with edge weights defined by Morisita’s clone overlap index, node colour denoting clonality, and node size denoting number of clones. For clarity of cell networks, we filtered out edges with very low clonal overlap. We defined these filter thresholds based on the distribution of Morisita’s index across all CD4^+^ and CD8^+^ T-cell populations, as these are likely to be spurious connections, considering all edges below the 95^th^ percentile as low confidence. Raw, unfiltered counts of all overlaps were visualized as heatmaps. Networks were visualized using the R package ggraph, with graphs laid out for visualization using a force-directed Fruchterman-Reingold layout.

## Statistics, data analysis, and visualisation

All statistics, data analysis, and visualisation was done in Python or R unless otherwise stated. Other single-cell analysis figures were produced using R packages including ggplot2, ggpubr, ComplexHeatmap (Gu et al., 2016); Python packages including muon (Bredikhin et al., 2022); and the Panpipes workflow (Curion et al., 2024). Significance is indicated as follows: \**P* < 0.05; \*\**P* < 0.01; \*\*\**P* < 0.001; \*\*\*\**P* < 0.0001. Schematics were produced using BioRender.com.

## Supplemental material

Supplementary Table 1. Demographics and study details of LEGACY study participants.

Supplementary Table 2. LEGACY cohort HLA genotypes and genotype-dependent ancestral composition.

Supplementary Figure 1. Detection of antigen-specific antibodies against influenza strains used in the 2022/23 influenza vaccine, and H1N1 and H5N8.

## Data availability

All raw and processed single-cell transcriptomics data will be available on CELLxGENE. Genotyping data will be available on GEO. All code will be available on Zenodo upon acceptance of this paper.

## Acknowledgments

We thank the participants and the PPIE members of the LEGACY committee for their support of this work. This research was conducted at the NIHR Imperial Clinical Research Facility, London UK, and supported by NIHR Imperial Biomedical Research Centre and the Single-Cell and Spatial Genomics Facility at the Kennedy Institute of Rheumatology, University of Oxford. JHS is supported by Wellcome Trust Early Career Award (226938/Z/23/Z). CAD is supported by the Wellcome Trust and Royal Society (204290/Z/16/Z), the UK Medical Research Council (MR/T030410/1), grant funding from Johnson & Johnson Innovative Medicine, and the NIHR Oxford Biomedical Research Centre, Inflammation Across Tissues and Cell and Gene Therapy Themes. KMP is supported by an UKRI/MRC Clinician Scientist Research Fellowship (MR/W024977/1). The study was funded by the Silicon Valley Community Foundation from a donation by the Chan Zuckerberg Initiative, the funder had no role in the study design, data analysis or decision to publish. The funder provided data infrastructure support for the posting of the online dataset with CELLxGENE.

## Author contributions

J.H.Y. Siu: Data curation, Formal analysis, Investigation, Methodology, Project Administration, Supervision, Visualization, Writing – original draft, Writing – review & editing; S. Coelho: Data curation, Investigation, Methodology, Project administration; A. Palomeras: Data curation, Methodology, Project administration; S. Belij-Rammerstorfer: Methodology, Investigation, Formal analysis; C.H. Lee: Investigation, Formal analysis; T. Strobel: Investigation, Formal analysis; C. Thorpe: Investigation, Formal analysis; C. Kaur: Data curation, Investigation; T. Cole: Project administration; N. Remmert: Formal analysis, Visualization; J. Fowler: Investigation; S. Pledger: Investigation; K.B. Dooley: Investigation; D. Opoka: Investigation; T. Szommer: Data curation, Project administration; S. Vanderslott: Funding acquisition, Methodology; P. Kaleebu: Conceptualization, Supervision, Funding acquisition; A. Milicic: Funding acquisition, Writing – review & editing; D.B. Palmer: Funding acquisition, Writing – review & editing; T. Lambe: Conceptualization, Funding acquisition, Resources, Supervision; B.D. Marsden: Data curation, Funding acquisition, Project administration, Supervision; H. Koohey: Funding acquisition, Supervision, Writing – review & editing; M. Coles: Conceptualization, Funding acquisition, Resources, Supervision, Writing – review & editing; C.A. Dendrou: Conceptualization, Formal analysis, Funding acquisition, Investigation, Methodology, Project administration, Resources, Supervision, Visualization, Writing – original draft, Writing – review & editing; K.M. Pollock: Conceptualization, Funding acquisition, Investigation, Methodology, Project administration, Resources, Supervision, Writing – original draft, Writing – review & editing.

