## Supplemental Materials for "Early regional lymph node activation drives influenza vaccine responses in an ancestrally diverse cohort"

**Contents:**

**Supplementary Table 1. Demographics and study details of LEGACY study participants.**

**Supplementary Table 2. LEGACY cohort HLA genotypes and genotype-dependent ancestral composition.**

**Supplementary Figure 1. Detection of antigen-specific antibodies against influenza strains used in the 2022/23 influenza vaccine, and H1N1 and H5N8.**

**Supplementary Table 1. Demographics and study details of LEGACY study participants.**

Study cohort demographics include age, sex, weight, height, body mass index (BMI), country of birth (standardised), parents' and grandparents' country of birth (standardised), and self-declared ethnicity (standardised to HANCESTRO ontology). Study details including day of post-vaccination FNA, number of days between LN FNAs, and vaccination arm. F, female; L, left; M, male; n.d., no data; R, right

| Subject ID | Age (yr) | Sex | Day of post-vaccination FNA | No. of days between FNAs | Vaccine arm | Weight (kg) | Height (m) | BMI | Country of birth | Parents' country of birth | | Grandparents' (GP) country of birth | | | | HANCESTRO ontology |
|---|---|---|---|---|---|---|---|---|---|---|---|---|---|---|---|---|
| | | | | | | | | | | Parent 1 | Parent 2 | GP1 | GP2 | GP3 | GP4 | |
| LEG1002 | 24 | F | 4 | 29 | L | 56.2 | 1.70 | 19.45 | India | India | India | India | India | India | India | Indian |
| LEG1004 | 50 | M | 5 | 27 | L | 79.7 | 1.87 | 22.79 | UK | Ireland | India | Ireland | Ireland | India | India | Indian/Irish |
| LEG1005 | 21 | F | 7 | 44 | L | 58.5 | 1.71 | 20.01 | China | China | China | China | China | China | China | Chinese |
| LEG1006 | 34 | F | 5 | 29 | R | 62.7 | 1.64 | 23.31 | France | Sri Lanka | Sri Lanka | Sri Lanka | Sri Lanka | Sri Lanka | Sri Lanka | Tamil |
| LEG1007 | 22 | F | 5 | 50 | L | 68.0 | 1.67 | 24.38 | China | China | China | China | China | China | China | Chinese |
| LEG1008 | 27 | F | 3 | 24 | R | 74.4 | 1.72 | 25.15 | UK | England | England | England | England | Jamaica | Jamaica | British/Afro-Caribbean |
| LEG1009 | 35 | M | 6 | 48 | R | 85.5 | 1.77 | 27.29 | UK | Bangladesh | Bangladesh | Bangladesh | Bangladesh | Bangladesh | Bangladesh | Bangladeshi |
| LEG1017 | 34 | F | 6 | 29 | L | 70.0 | 1.59 | 27.69 | UK | Nigeria | Nigeria | Nigeria | Nigeria | Nigeria | Nigeria | Yoruban |
| LEG1018 | 38 | M | 7 | 42 | L | 61.0 | 1.59 | 24.13 | UK | India | Kenya | India | India | India | India | Indian |
| LEG1019 | 41 | F | 5 | 33 | L | 52.1 | 1.64 | 19.37 | UK | England | Nigeria | England | England | Nigeria | Nigeria | British/ Yoruban |
| LEG1021 | 40 | M | 6 | 13 | L | 89.0 | 1.78 | 28.09 | India | India | India | India | India | India | India | Indian |
| LEG1025 | 32 | M | 6 | 14 | L | 93.8 | 1.86 | 27.11 | UK | Egypt | Egypt | Egypt | Egypt | n.d. | n.d. | Arab |
| LEG1028 | 19 | F | 6 | 13 | L | 53.4 | 1.60 | 20.86 | UK | Philippines | Philippines | Philippines | Philippines | Philippines | Philippines | Filipino |

**Supplementary Table 2. LEGACY cohort HLA genotypes and genotype-dependent ancestral composition.**

Participant HLA genotyping (up to four digits) and ancestry as determined by genotype-dependent admixture analysis (at K=18). AFR, African; AMR, Admixed American; CSA, Central/South Asian; EAS, East Asian; EUR, European; MDE, Middle Eastern; n.f., not found; OCE, Oceanian; SAS, South Asian.

| Subject ID | HLA-A* allele 1 | HLA-A* allele 2 | HLA-B* allele 1 | HLA-B* allele 2 | HLA-C* allele 1 | HLA-C* allele 2 | HLA-DRB1* allele 1 | HLA-DRB1* allele 2 | HLA-DRB3/4/5* allele 1 | HLA-DRB3/4/5* allele 2 | HLA-DQB1* allele 1 | HLA-DQB1* allele 2 | AMR | OCE | CSA | AFR | MDE | EAS | EUR | SAS |
|---|---|---|---|---|---|---|---|---|---|---|---|---|---|---|---|---|---|---|---|---|
| LEG1002 | 24:02 | 31:01 | 37:01 | 40:01 | 03:04 | 06:02 | 04:04 | 10:01 | DRB4 | DRB4 | 03:02 | 05:01 | 0.018 | 0.009 | 0.220 | 0.009 | 1.00 E-05 | 0.012 | 0.096 | 0.637 |
| LEG1004 | 02:01 | 33:03 | 08:01 | 44:03 | 07:01 | 07:01 | 03:01 | 07:01 | DRB3 | DRB4 | 02 | 02 | 0.009 | 0.009 | 0.149 | 0.002 | 1.00 E-05 | 0.019 | 0.565 | 0.246 |
| LEG1005 | 24:02 | 33:03 | 39:01 | 58:01 | 03:02 | 03:04 | 13:02 | 15:01 | DRB3 | DRB5 | 06:02 | 06:09 | 0.009 | 1.00 E-05 | 2.00 E-05 | 4.00 E-05 | 1.00 E-05 | 0.991 | 2.00 E-05 | 1.00 E-05 |
| LEG1006 | 11:01 | 24:02 | 35:01 | 44:03 | 04:01 | 04:01 | 07:01 | 15:02 | DRB4 | DRB5 | 02 | 06:01 | 0.008 | 0.004 | 0.061 | 0.008 | 0.034 | 0.047 | 0.020 | 0.817 |
| LEG1007 | 11:01 | 33:03 | 40:01 | 58:01 | 03:02 | 07:02 | 03:01 | 04:04 | DRB3 | DRB4 | 02 | 03:02 | 0.014 | 1.00 E-05 | 2.00 E-05 | 4.00 E-05 | 1.00 E-05 | 0.986 | 2.00 E-05 | 1.00 E-05 |
| LEG1008 | 03:01 | 74:01 | 07:02 | 15:03 | 02:10 | 07:02 | 01:01 | 11:01 | DRB3 | n.f. | 03:01 | 05:01 | 0.004 | 0.005 | 0.044 | 0.504 | 1.00 E-05 | 4.00 E-05 | 0.443 | 1.00 E-05 |
| LEG1009 | 32:01 | 33:03 | 44:03 | 49:01 | 07:01 | 07:01 | 07:01 | 11:03 | DRB3 | DRB4 | 02 | 03:01 | 0.028 | 0.010 | 0.064 | 0.001 | 0.016 | 0.133 | 0.099 | 0.649 |
| LEG1017 | 01:01/04N | 34:02 | 58:01 | 58:01 | 07:01 | 16:01 | 15:03 | 15:03 | DRB5 | DRB5 | 06:02 | 06:02 | 3.00 E-05 | 3.49 E-04 | 2.00 E-05 | 0.100 | 1.00 E-05 | 4.00 E-05 | 2.00 E-05 | 1.00 E-05 |
| LEG1018 | 01:01/04N | 23:01 | 40:06 | 44:03 | 04:01 | 15:02 | 07:01 | 11:01 | DRB3 | DRB4 | 02 | 03:01 | 0.028 | 0.030 | 0.108 | 0.007 | 0.032 | 0.011 | 0.037 | 0.746 |
| LEG1019 | 01:01/04N | 03:01 | 44:02 | 57:04 | 05:01/03 | 18:01 | 04:01 | 15:01 | DRB4 | DRB5 | 02 | 03:01 | 0.010 | 1.00 E-05 | 0.067 | 0.510 | 0.072 | 4.00 E-05 | 0.341 | 1.00 E-05 |
| LEG1021 | 03:01 | 32:01 | 35:03 | 52:01 | 12:02 | 12:03 | 09:01 | 15:02 | DRB4 | DRB5 | 03:03 | 06:01 | 0.019 | 0.010 | 0.215 | 0.003 | 1.00 E-05 | 0.002 | 0.062 | 0.689 |
| LEG1025 | 03:02 | 23:01 | 08:01 | 14:02 | 07:02 | 08:02 | 03:01 | 03:02 | DRB3 | DRB3 | 02 | 04:02 | 0.003 | 1.00 E-05 | 0.162 | 0.193 | 0.413 | 0.012 | 0.217 | 1.00 E-05 |
| LEG1028 | 11:01 | 24:02 | 38:01 | 40:10 | C04:03 | 12:03 | 13:02 | 15:01 | DRB3 | DRB5 | 05:02 | 06:03 | 0.016 | 0.034 | 2.00 E-05 | 0.010 | 0.013 | 0.827 | 0.091 | 0.006 |

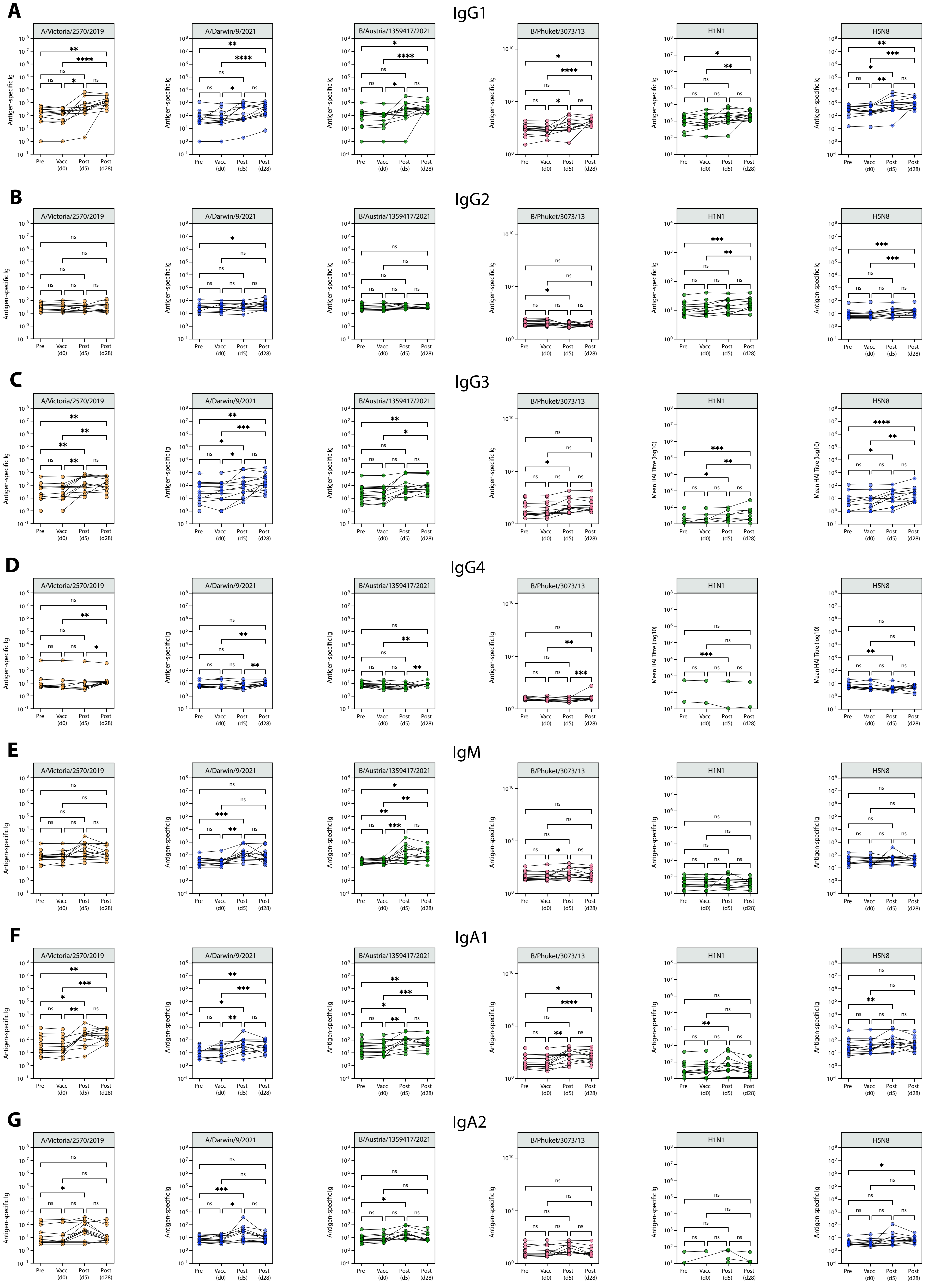


**Supplementary Figure 1. Detection of antigen-specific antibodies against influenza strains used in the 2022/23 influenza vaccine, and H1N1 and H5N8.**

Detection of antigen-specific antibodies - **(A)** IgG1, **(B)** IgG2, **(C)** IgG3, **(D)** IgG4, **(E)** IgM, **(F)** IgA1 and **(G)** IgA2 - by Luminex assay at various timepoints (pre: pre-vaccination, vacc: day of vaccination, post (d5): median five days post-vaccination, post (d28): median 28 days post-vaccination).
